# Phosphorylation-dependent regulation of Hsp70 chaperones stimulates client recruitment

**DOI:** 10.64898/2026.06.01.729190

**Authors:** Yun Chen, Yu-Yen Wang, Yun-Li Yew, Yuan-Teng Chang, Ting-Yu Liu, Shu-Chun Teng

## Abstract

Molecular chaperones of the Hsp70 family play essential roles in maintaining proteostasis, particularly under conditions of cellular stress. Posttranslational modifications of Hsp70, collectively termed the chaperone code, are emerging as critical regulators of chaperone function, yet their mechanistic contributions remain incompletely understood. Here, we investigate the functional significance of a conserved phosphorylation site in Hsp70, corresponding to serine 326 in yeast Ssa1 and serine 329 in human HSPA8. We demonstrate that DNA double-strand break stress increases cellular reliance on Hsp70 activity in yeast, highlighting a role for chaperones in the DNA damage response. Loss of Ssa1 serine 326 phosphorylation impairs multiple Hsp70-dependent functions, including prion sequestration and glucocorticoid receptor maturation, indicating that this modification is required for optimal chaperone activity. Extending these findings to human cells, we show that the homologous HSPA8 serine 329 residue is necessary for efficient clearance of polyglutamine aggregates. Mechanistically, a phospho-deficient HSPA8-S329A mutant exhibits reduced client-binding capacity. Together, our findings identify a conserved phosphorylation event that enhances Hsp70 function by promoting client engagement, providing new insight into how the chaperone code regulates proteostasis across species.

## Introduction

Protein homeostasis is essential for cellular function and survival. Under environmental stress conditions, including DNA damage, oxidative stress, and heat shock, the cellular protein-folding environment becomes destabilized, leading to protein misfolding and the accumulation of toxic aggregates [1, 2]. Heat shock proteins (HSPs), which are highly conserved across prokaryotes and eukaryotes, comprise a major class of molecular chaperones that safeguard proteome integrity [3–5]. These chaperones coordinate multiple quality control processes, including nascent protein folding, refolding of misfolded species, and the clearance or disassembly of protein aggregates [5–7].

Hsp70 proteins are highly conserved from yeast to humans, with Ssa1 and Ssa2 serving as the major cytosolic Hsp70s in *Saccharomyces cerevisiae*, and their constitutively expressed human homolog being HSPA8 (also known as HSC70) [8]. Hsp70 molecular chaperones function through an ATP-dependent chaperone cycle, in which co-chaperones, particularly J-domain proteins, initially recognize unfolded or misfolded client proteins and deliver them to Hsp70 for refolding [9]. This process is further regulated by nucleotide exchange factors (NEF), which modulate ATP hydrolysis and control substrate binding and release [10].

Accumulating evidence suggests that the DNA damage response (DDR) kinases ATM and ATR function not only in genome surveillance but also in the regulation of protein homeostasis. During DNA metabolism, cells continuously encounter diverse forms of genomic damage and therefore rely on sophisticated DDR networks to maintain genome integrity [13,14]. ATM and ATR act as master transducers of DNA damage signals [15,16], phosphorylating numerous downstream substrates, primarily at SQ/TQ motifs, to coordinate DNA repair, cell cycle checkpoints, and cellular adaptation to stress [15,17,18]. Beyond these canonical functions, ATM and ATR have also been implicated in proteostasis regulation. In mammalian systems, ATM and ATR contribute to neuronal vesicle dynamics [11], while defects in ATM-mediated oxidative stress responses are associated with increased proteotoxic stress and widespread protein aggregation [2]. Similarly, disruption of the yeast DDR kinase Mec1 exacerbates polyQ aggregation [12], suggesting that the role of DDR kinases in proteostasis is evolutionarily conserved. Consistently, loss of ATM or ATR activity promotes protein aggregation in both yeast and human cells [12], indicating that DDR signaling is critical for maintaining proteostasis under genotoxic stress. Together, these findings highlight extensive crosstalk between genome stability pathways and proteostasis networks; however, the molecular mechanisms linking DNA damage signaling to chaperone-mediated protein quality control remain poorly understood.

To explore whether Hsp70 may serve as a regulatory node linking these processes, in this study, we examined whether phosphorylation at conserved residues constitutes an additional regulatory mechanism controlling Hsp70 function during cellular stress. We compiled all potential ATM/ATR phosphorylation sites containing SQ/TQ motifs within yeast Hsp70 proteins and annotated previously reported phosphorylation events based on available databases. This analysis revealed that the cytosolic Hsp70 Ssa1 S326 in yeast is subject to phosphorylation [19]. Notably, this residue is highly conserved across eukaryotic species, corresponding to serine 326 in yeast Ssa1 and serine 329 in human HSPA8, suggesting a conserved regulatory mechanism. We demonstrate that phosphorylation of Hsp70 at S326 is required for cellular survival under DNA double-strand break (DSB) stress and is critical for maintaining optimal chaperone activity. Furthermore, we show that the corresponding residue in human HSPA8 facilitates the resolution of Huntington’s disease-related HTT-polyQ aggregation by enhancing its interaction with polyQ substrates. Together, our findings identify a conserved, DNA damage-associated phosphorylation event on Hsp70 that modulates chaperone activity and proteostasis.

## Results

### Hsp70 phosphorylation is essential for yeast survival following DNA double-strand breaks (DSBs)

The yeast *Saccharomyces cerevisiae* contains two constitutively expressed Hsp70 proteins, Ssa1 and Ssa2. To explore the functional importance of Hsp70 under different stress conditions, yeast spotting assays were performed using wild-type (WT), *ssa1Δ*, *ssa2Δ*, and *ssa1Δ ssa2Δ* strains exposed to several cellular stresses, including heat shock, methyl methanesulfonate (MMS) (DNA alkylation stress), bleomycin (DNA DSBs), KCl (osmotic stress), H₂O₂ (oxidative stress), and rapamycin (translation inhibition). As expected from previous reports, the *ssa1Δ ssa2Δ* double mutant exhibited a severe growth defect under heat stress (39 °C) [20]. In addition to this known phenotype, the double mutant also displayed increased sensitivity to DNA-damaging agents, including MMS and bleomycin. In contrast, deletion of *SSA1* or *SSA2* alone did not result in overt growth defects under other tested conditions (Figure 1A). These results indicate that yeast Hsp70 activity becomes particularly important for cellular survival following induction of DNA damage.

**Figure 1.**
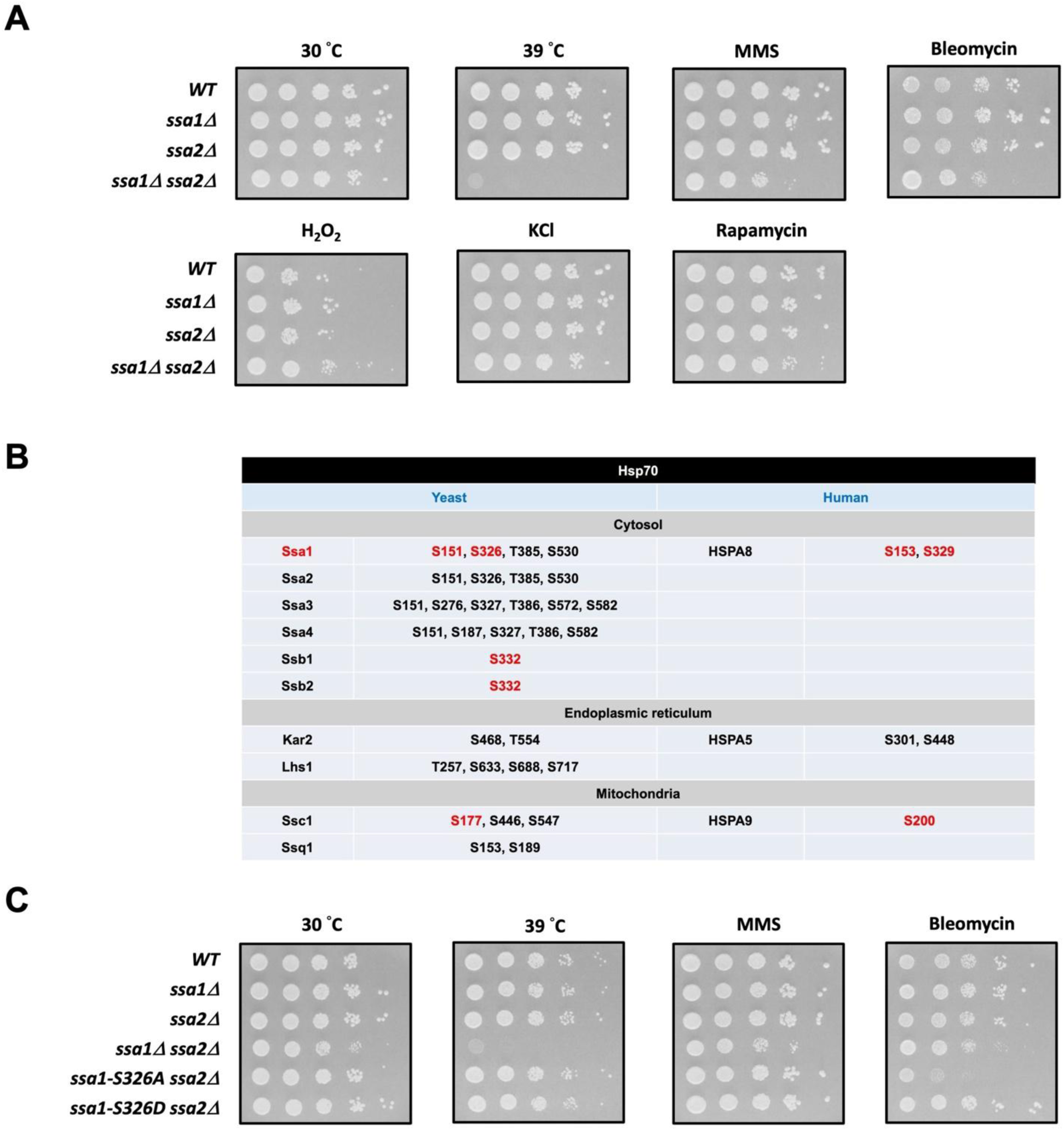
Yeast Hsp70 function is required for growth under DNA DSB stress. (A) The WT, *ssa1Δ*, *ssa2Δ*, and *ssa1Δ ssa2Δ* cells were grown to log phase and spotted in 10-fold serial dilution on SC stress-containing agar plates as indicated. (B) Multiple SQ/TQ motifs were identified across yeast cytosolic Hsp70 proteins, with previously validated phosphorylation sites marked in red based on the database. (C) The WT, *ssa1Δ*, *ssa2Δ*, *ssa1Δ ssa2Δ*, *ssa1-S326A ssa2Δ*, and *ssa1-S326D ssa2Δ* cells were grown to log phase and spotted in 10-fold serial dilution on SC stress-containing agar plates.

To understand whether yeast Hsp70 proteins may be regulated in response to DNA damage, we systematically examined all yeast Hsp70 family members for the presence of conserved SQ/TQ motifs, which are commonly associated with DNA damage-responsive phosphorylation events. Multiple SQ/TQ sites were identified across yeast Hsp70 proteins, and several of these sites have been previously detected as phosphorylated residues by mass spectrometry according to the BioGRID database (Figure 1B). Given that serine 326 of Ssa1 resides within a conserved SQ motif and represents a candidate DNA damage-responsive phosphorylation site, we next examined whether the phosphorylation state of this residue contributes to the cellular response to DNA damage. Phospho-deficient (*ssa1-S326A ssa2Δ*) and phospho-mimetic (*ssa1-S326D ssa2Δ*) strains were generated and analyzed under bleomycin and MMS treatment. While MMS treatment did not cause noticeable growth defects in either mutant strain, cells expressing the Ssa1-S326A mutant exhibited markedly reduced growth in the presence of DNA bleomycin, resembling the phenotype observed in the *ssa1Δ ssa2Δ* strain. In contrast, expression of the Ssa1-S326D mutant substantially rescued growth under bleomycin treatment (Figure 1C). These findings suggest that phosphorylation of Ssa1 at Ser326 plays an important role in supporting Hsp70 function during the response to DNA DSBs.

### Loss of Ssa1 serine 326 phosphorylation impairs Hsp70 chaperone functions

To assess whether phosphorylation of Ssa1 at serine 326 influences Hsp70 function, we analyzed prion propagation in yeast. Prion propagation was first assessed using the Sup35-based *[PSI⁺]* reporter system, in which efficient chaperone activity promotes sequestration of Sup35 into prion aggregates, resulting in suppression of translation termination and the appearance of white colonies. In contrast, impaired chaperone function favors soluble Sup35, leading to efficient translation termination and red colony formation. Under these conditions, the *ssa1-S326A* strain displayed a pronounced shift toward red colony coloration, indicating defective prion propagation compared with strains expressing wild-type or *ssa1-S326D* (Figure 2A). These results suggest that the phospho-deficient *ssa1-S326A* mutant is compromised in supporting Hsp70-dependent prion maintenance.

**Figure 2.**
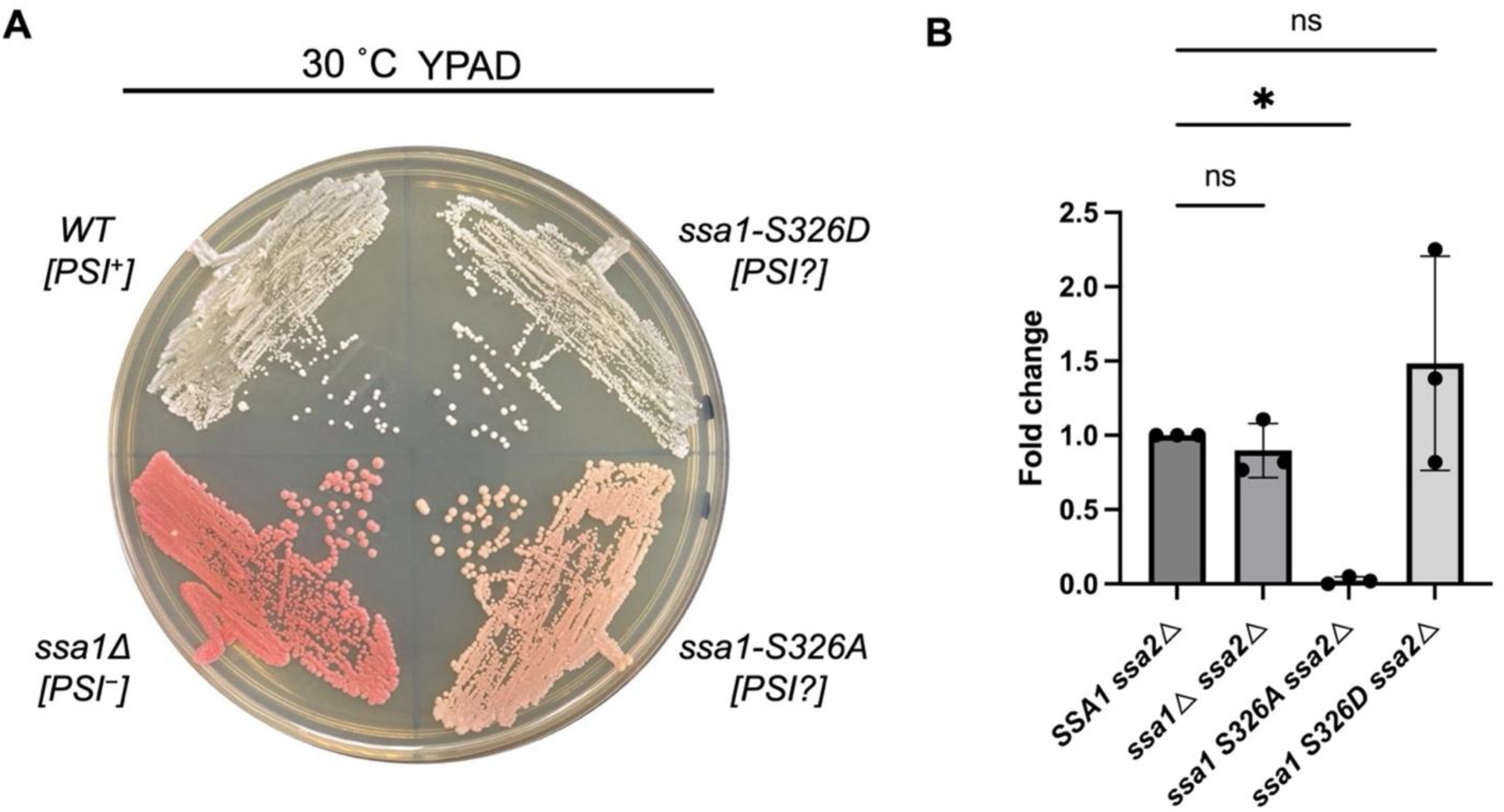
Ssa1 S326 modifications affect prion propagation and glucocorticoid receptor (GR) activation. (A) WT, *ssa1Δ*, *ssa1-S326A*, and *ssa1-S326D* 5V-H19 yeast strains were streaked on the YEPD agar plate. The Sup35 prion protein in the 5V-H19 yeast strain can frequently read through the stop codon of the mRNA from the *ade2-1* allele. (B) The W303 WT, *SSA1 ssa2*, *ssa1, ssa2*, *ssa1-S326A ssa2*, and *ssa1-S326D ssa2* cells were transformed with the GR expression vector (pG/N795) and the reporter plasmid with GR response elements (pUCΔSS-26X). Single clones were grown at 30°C in minimal medium to log phase. The relative GR activity is the mean of three independent experiments. Error bars indicate standard errors.

We next evaluated Hsp70 chaperone function using a GR activation assay, which monitors the ability of Hsp70 to assist the folding and maturation of newly synthesized GR into a competent state. Consistent with the prion propagation results, GR activity was nearly abolished in *ssa1-S326A ssa2Δ* cells, whereas robust GR activation was observed in cells expressing wild-type or *ssa1-S326D* (Figure 2B). Together, these findings demonstrate that loss of Ser326 phosphorylation severely impairs core Hsp70 chaperone functions, affecting both protein quality control and client protein activation.

### HSPA8 serine 329, the human homolog of yeast Ssa1 serine 326, is required for efficient clearance of polyQ aggregates

To investigate whether the conserved phosphorylation site corresponding to yeast Ssa1 Ser326 regulates Hsp70 function in human cells, we first performed a sequence alignment and found that Ssa1 serine 326 is highly conserved across eukaryotes. The human homolog, HSPA8, contains the corresponding residue serine 329 near the end of the nucleotide-binding domain (NBD) (Figure 3A).

**Figure 3.**
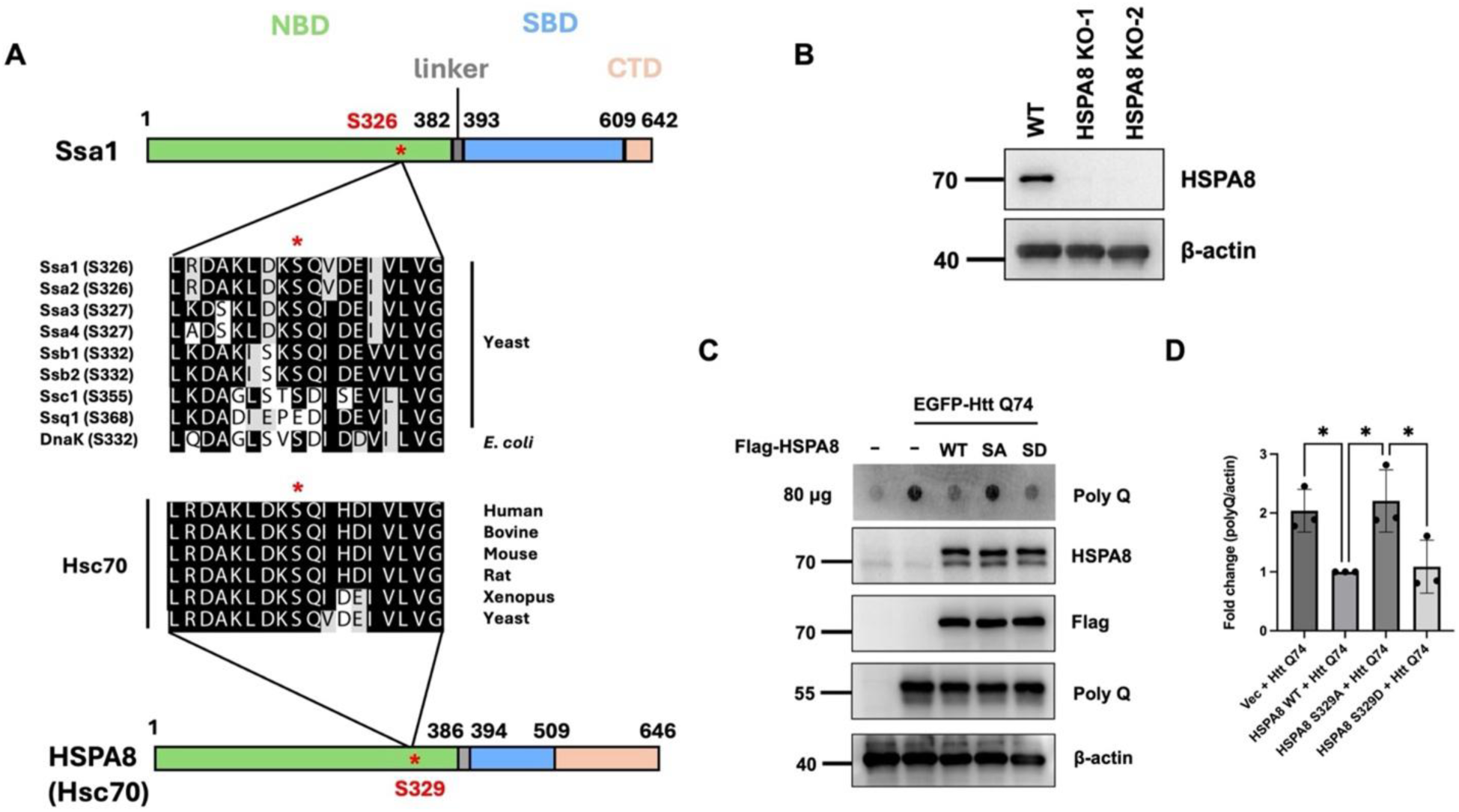
Phosphorylation of human HSPA8 S329, corresponding to yeast Ssa1 S326, is required for clearance of polyQ aggregates. (A) The domain architecture of Ssa1 is shown, with S326 indicated by an asterisk (*). Sequence alignment surrounding Ssa1 S326 was performed among yeast Hsp70 family members, *E. coli* DnaK, and Hsp70 homologs from diverse animal species. (B) Generation of HSPA8 KO HeLa cells using CRISPR/Cas9. (C) Representative chemiluminescent images of Htt polyQ, HSPA8 (WT and S329 mutants), and β-actin are shown. Dephospho-mimetic (SA) and phospho-mimetic (SD) variants were generated by site-directed mutagenesis at the indicated residue. HSPA8 KO HeLa cells were first transfected with HSPA8 constructs for 24 h, followed by transfection with Htt polyQ for an additional 48 h. PolyQ aggregation was analyzed by the filter trap assay. Data were analyzed by one-way ANOVA followed by Tukey’s post hoc test. (D) Quantification of polyQ aggregation, normalized to β-actin, in HSPA8 knockout HeLa cells expressing Flag-tagged HSPA8 WT or S329 variants. Statistical significance was indicated by *, P < 0.05.

To study the functional relevance of HSPA8 Ser329, we generated HSPA8 knockout (KO) HeLa cells using CRISPR/Cas9 (Figure 3B). These cells were subsequently transfected with HSPA8 wild-type, phospho-deficient (S329A), or phospho-mimetic (S329D) expressing plasmid. Upon expression, cells harboring HSPA8-S329A exhibited a marked impairment in resolving polyQ-expanded mutant huntingtin (Htt) aggregates compared with those expressing wild-type or S329D HSPA8 (Figure 3C and 3D). These results indicate that phosphorylation of HSPA8 at serine 329 is critical for its ability to maintain proteostasis of aggregation-prone Htt client proteins in human cells.

### Phospho-deficient HSPA8-S329A exhibits reduced client binding

To investigate the molecular basis for the impaired clearance of Htt polyQ aggregates by HSPA8-S329A, co-immunoprecipitation (Co-IP) assays were performed to assess the interaction between HSPA8 and its client proteins. Compared with wild-type or HSPA8-S329D, the S329A mutant displayed substantially reduced binding to its substrate (Figure 4A and 4B). These results indicate that phosphorylation at Ser329 is critical for maintaining HSPA8’s ability to engage and chaperone client proteins, providing a mechanistic explanation for the defective polyQ aggregate resolution observed in S329A-expressing cells (Figure 3C).

**Figure 4.**
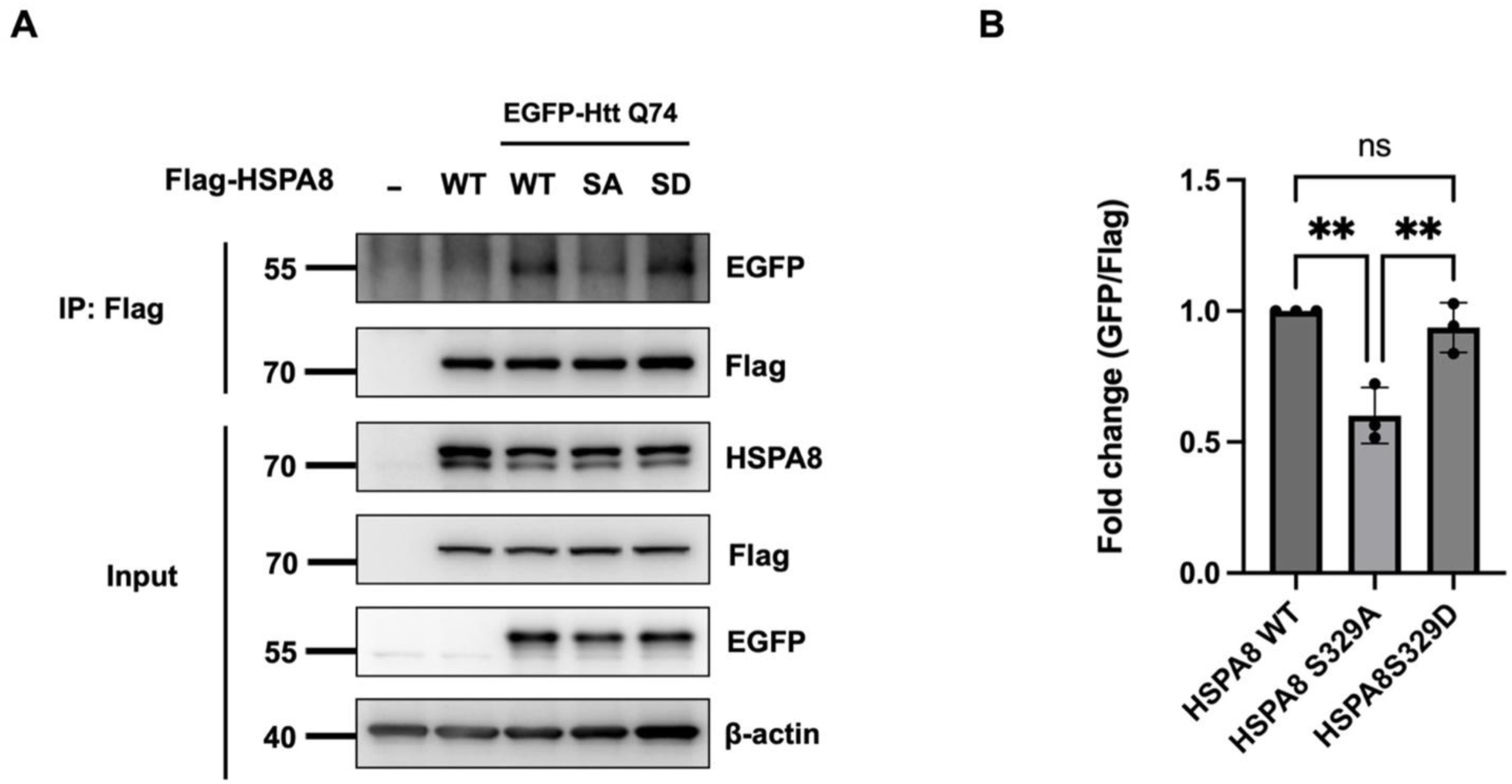
Phospho-deficient HSPA8-S329A exhibits reduced client binding. (A) Lysates from HSPA8 knockout HeLa cells expressing Flag-tagged HSPA8 WT or S329 mutants were immunoprecipitated with anti-Flag antibodies, followed by immunoblotting to detect co-precipitated EGFP-tagged Htt polyQ. (B) Comparison of the interaction of EGFP–Htt polyQ with Flag-tagged HSPA8 WT and S329 mutants. Statistical significance was indicated by **, P < 0.01.

**Figure 5.**
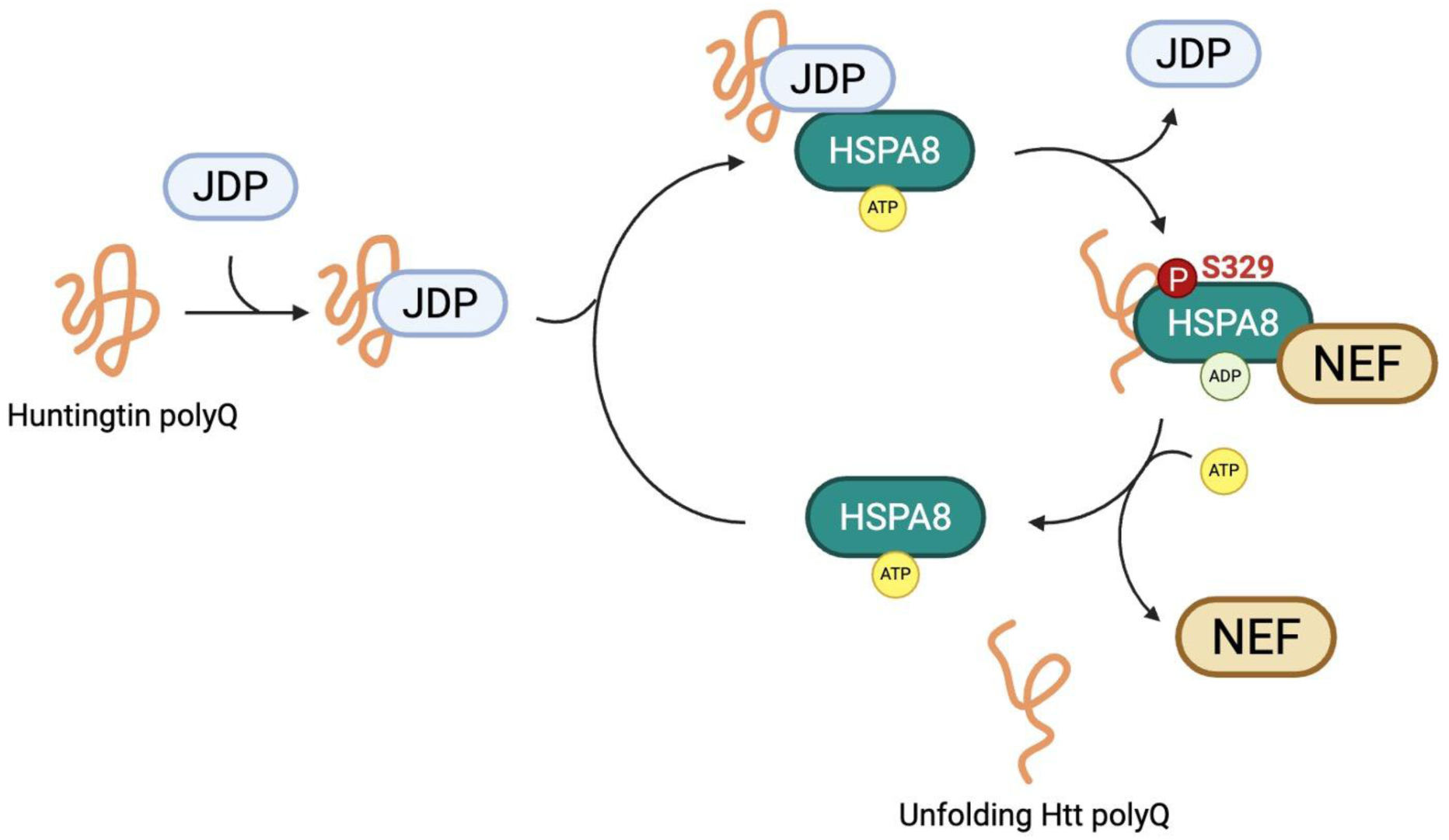
Proposed model for the role of HSPA8 serine 329 phosphorylation in regulating Hsp70 chaperone activity. HSPA8 mediates protein folding through an ATP-dependent chaperone cycle. In the ATP-bound state, HSPA8 interacts with J-domain proteins (JDPs) and client Htt polyQ proteins. ATP hydrolysis leads to a high-affinity state that stabilizes client binding. Nucleotide exchange by NEFs promotes ADP release and restores the ATP-bound state, allowing substrate release and completion of the folding cycle. Phosphorylation of HSPA8 at serine 329 enhances its association with Htt PolyQ, thereby promoting its proper folding and facilitating its degradation.

## Discussion

In this study, we identify a conserved phosphorylation site within Hsp70 that plays a critical role in maintaining chaperone function under stress conditions. Using yeast as a model system, we reveal that Hsp70 activity becomes particularly important for cell survival in response to DNA DSB stress and that mutation of Ssa1 S326 compromises this function. Functional assays further demonstrate that disruption of this site impairs core Hsp70-dependent processes, including prion propagation and client protein activation/maturation. Extending these findings to human cells, we found that the phosphorylation state of the corresponding residue, HSPA8 S329, regulates the clearance of aggregation-prone proteins such as Htt polyQ. The phospho-deficient HSPA8-S329A mutant also shows reduced client Htt polyQ binding, suggesting a possible mechanism for its impaired chaperone activity. Together, our results reveal a conserved regulatory mechanism that links Hsp70 function to cellular stress responses and proteostasis.

In our study, phosphorylation of Ssa1 at S326 appears to be particularly important under conditions of DNA DSB. This effect could reflect a reduced capacity of Ssa1-S326A to assist the folding or stabilization of certain client proteins that participate in DNA DSB repair. Previous studies indicate that heat shock proteins, particularly Hsp70 and Hsp90, play an active role in maintaining genome stability [21]. They are not only responsible for rescuing misfolded proteins but also directly participate in the sensing, signal transduction, and effector steps of the DNA damage response [21]. Through these activities, HSPs support the proper function of multiple DNA repair proteins, reinforcing their contribution to cellular survival under DNA-damaging conditions. Together, these observations suggest that the phosphorylation state of Ssa1 at S326 may influence yeast cell fitness during DNA damage, potentially by affecting the maturation or activity of DNA DSB repair-related client proteins.

Hsp70 function is commonly divided into two major domains. The NBD regulates ATP binding and hydrolysis and interacts with NEFs [10]. In contrast, the substrate-binding domain (SBD) is responsible for binding client proteins [22]. Although these domains have distinct roles, they are closely connected through dynamic and bidirectional communication [23, 24]. ATP binding and hydrolysis in the NBD drive changes in protein shape that control the opening and closing of the SBD, thereby regulating how tightly substrates/clients are bound [23, 25]. In turn, substrate binding to the SBD can stimulate ATP hydrolysis in the NBD, forming a coordinated cycle that is essential for Hsp70 function [23]. In this context, S329 is located near the NBD-linker junction, a region important for communication between the two domains [23]. This suggests that phosphorylation at this site may be more likely to affect domain coordination rather than directly altering substrate binding. Consistent with this idea, the reduced substrate association observed in the HSPA8-S329A mutant may result from a decreased ability to switch between functional states required for efficient client binding. In addition, phosphorylation at this site may influence interactions with co-chaperones [26]. Alternatively, changes in protein shape may also affect how Hsp70 supports proper protein folding. These possibilities warrant further investigation.

## Methods

### Yeast strains and plasmids

All yeast strains used in this study were derived from W303 (*MATa/α ura3-52/ura3-52 trp1Δ2/ trp1Δ 2 leu2-3_112/ leu2-3_112 his3-11/ his3-11 ade2-1 can1-100/ade2-1 can1-100*) and 5V-H19 (*SUQ5 ade2-1(UAA) can1-100 leu2-3,112 ura3-52*). The single gene deletion strains were obtained from the deletion library (Invitrogen), while the double gene deletion strains were selected by tetrad dissection. Standard genetic and cloning methods were used for all constructions.

### Yeast growth media

The yeast media used were rich medium (YEP, 1% yeast extract, 2% peptone) or synthetic complete (SC) medium containing 2% glucose.

### Spotting assay

W303-derived yeast strains (*ssa1Δ, ssa2Δ, ssa1Δ ssa2Δ, ssa1-S326A ssa2Δ, and ssa1-S326D ssa2Δ*) were grown to log phase, serially diluted (10-fold), and spotted on SC agar plates containing the indicated stress conditions (15 mU/mL Bleomycin, 0.005% MMS, 2 mM H_2_O_2_, 0.5 M KCl, 150 ng/mL Rapamycin). Plates were incubated at 30℃ or 39℃ as indicated.

### Yeast GR activity assay (lacZ reporter assay)

W303-derived yeast strains were co-transformed with the glucocorticoid receptor (GR) expression vector (pG/N795) and a GR-responsive reporter plasmid (pUCΔSS-26X), and maintained in SC-Ura-Trp medium at 30°C. Log-phase cultures were treated with 10 μM deoxycorticosterone (DOC) or DMSO control for 5 h. Cells were lysed in Z buffer (60 mM Na₂HPO₄, 60 mM NaH₂PO₄, 10 mM KCl, 1 mM MgSO₄, and 50 mM β-mercaptoethanol) with chloroform (1:3, v/v) and 0.01% sodium dodecyl sulfate (SDS), followed by vortexing with glass beads for 1 min. β-galactosidase activity was measured using 1 mg/mL o-nitrophenyl-β-D-galactopyranoside (ONPG) at 30°C for 10 min and stopped by the addition of 1 M Na₂CO₃. Absorbance at 420 nm was recorded. Data represent three independent biological replicates and were analyzed by one-way ANOVA.

### Human cell culture

HSPA8 knockout HeLa cells were cultured in high-glucose Dulbecco’s modified Eagle’s medium (DMEM) (Cytiva, Logan, UT, USA) supplemented with 10% fetal bovine serum (FBS) and 1× penicillin/streptomycin/fungizone. Cells were maintained at 37 °C in a humidified incubator with 5% CO₂ and were routinely screened to confirm the absence of mycoplasma contamination.

### Plasmids construction

pRS306*-SSA1* was constructed by polymerase chain reaction (PCR) amplifying a DNA fragment containing the full open reading frame and the downstream 300 nt from *S. cerevisiae* genomic DNA and ligating it into the pRS306 vector. pRS306*-ssa1-S326A* and pRS306*-ssa1-S326D* were generated by site-directed mutagenesis. To generate *SSA1* mutants, the pRS306*-ssa1* point-mutation plasmids were linearized with SalI (New England Biolabs) and transformed into yeast cells. *URA3* pop-out mutants were selected on 5-fluoroorotic acid (5-FOA)-containing plates. The *SSA1* mutations were confirmed by PCR and sequencing. The site-directed mutagenesis oligo sequences are provided in the Reagents and tools table.

pG/N795 (GR expression) and pUCΔSS-26X (GR reporter) were purchased from Addgene #108220 and #108218. p-FLAG-CMV2-HSPA8 was contructed by amplifyling HSPA8 from 293T cDNA and cloned into p-FLAG-CMV2 vector. p-FLAG-CMV2-HSPA8 S329A and S329D were generated by site-directed mutagenesis. The site-directed mutagenesis oligo sequences are provided in the Reagents and tools table. pEGFP-Q74 was acquired from Addgene #40262.

### Generation of HSPA8 KO HeLa cell lines

HSPA8 KO HeLa cells were generated using a CRISPR/Cas9-based genome editing approach. Single-guide RNAs (sgRNAs) targeting exon 2 of the human *HSPA8* gene were designed using the CRISPR Guide RNA Design Tool (Benchling). Two independent sgRNA sequences were cloned into the pSpCas9(BB)-2A-Puro (PX459) V2.0 expression vector. The resulting CRISPR-sgRNA constructs were transfected into HeLa cells using Non-liposome Transfection Reagent II (T-Pro NTR II). Following transfection, cells were subjected to puromycin selection (1 μg/mL) for 2 days. Surviving cells were isolated as single-cell clones and subsequently expanded for downstream analyses. The sgRNA sequences and primer information used for clone validation are provided in the Reagents and tools table.

### Transfection

HSPA8 knockout HeLa cells (2 × 10⁶) were plated in 10-cm culture dishes and allowed to adhere for 24 h before transfection. Plasmid DNA (2 μg) was transfected into cells using T-Pro Non-liposome Transfection Reagent II at a DNA-to-reagent ratio of 1:2 in the presence of 500 μL OPTI-MEM. The mixture of DNA and T-Pro NTR II Transfection Reagent was then incubated at room temperature for 15 min. Following brief centrifugation, the transfection mixture was added directly to the culture medium without antibiotics.

### Filter trap assay

For preparation of lysates, cultured cells were first collected and resuspended in filter trap lysis buffer containing 1% Triton X-100 in 1× PBS (pH 7.4), supplemented with 1 mM PMSF and Complete EDTA-free protease inhibitor cocktail (Roche, Basel, Switzerland). The suspension was subjected to brief sonication to ensure efficient disruption and homogenization. Protein amounts in each sample were quantified using the Bio-Rad Protein Assay (Bio-Rad Laboratories, Hercules, CA, USA). For the filter trap assay, lysates were diluted with lysis buffer containing 1% SDS to reach a final protein concentration of 1 μg/mL. A 0.2-μm nitrocellulose membrane was pretreated in rinse buffer composed of 1% SDS in 1× PBS (pH 7.4) and assembled in a 96-well dot-blot filtration apparatus (Bio-Rad Laboratories). The diluted samples were loaded onto the membrane and vacuum-filtered. Following filtration, the membrane was washed once with TBST buffer (150 mM NaCl, 20 mM Tris-HCl, pH 7.4, 0.05% Tween-20). Detection of trapped proteins was performed by immunostaining.

### Western blotting

Total cellular proteins were isolated and separated by SDS-polyacrylamide gel electrophoresis (SDS-PAGE), followed by electrotransfer onto a polyvinylidene difluoride (PVDF) membrane (Millipore). The membrane was incubated in 5% nonfat milk at room temperature to block nonspecific binding. Subsequently, the membrane was incubated with the indicated primary antibodies at 4 °C overnight; detailed antibody information is provided in the Reagents and tools table. After primary antibody incubation, the membrane was rinsed three times with TBST buffer (150 mM NaCl, 20 mM Tris-HCl, pH 7.4, 0.05% Tween-20). The membrane was then incubated with horseradish peroxidase-conjugated anti-mouse or anti-rabbit secondary antibodies (Jackson ImmunoResearch Inc., West Grove, PA, USA). Immunoreactive signals were visualized using Luminata™ Crescendo Western HRP Substrate (Millipore).

### Co-IP

HSPA8 knockout HeLa cells (2 × 10⁶) were seeded in 10-cm dishes and cultured for 24 h before transfection. Cells were first transfected with 3 μg of pFlag-HSPA8 plasmid for 24 h, followed by a second transfection with 1 μg of Htt polyQ-GFP plasmid for an additional 48 h. Cells were then harvested and lysed in immunoprecipitation (IP) buffer containing 150 mM NaCl, 0.1% NP-40, 1 mM EDTA, 1 mM PMSF, and 50 mM Tris-HCl (pH 7.5), supplemented with Complete EDTA-free protease inhibitor cocktail and PhosSTOP phosphatase inhibitors (Roche). For immunoprecipitation, 500 μg of total lysate in a final volume of 500 μL was incubated with 2 μg of anti-Flag antibody (F3165, Sigma) for 2 h at 4 °C with gentle rotation. Immune complexes were captured by incubation with protein G Mag Sepharose Xtra magnetic beads (Cytiva) for 1 h at 4 °C. Beads were washed three times with IP buffer to remove nonspecific binding. Bound proteins were eluted in 20 μL of 2× SDS sample buffer (250 mM Tris-HCl pH 6.8, 10% SDS, 0.25% bromophenol blue, 50% sucrose, 0.5 M 2-mercaptoethanol) and subsequently analyzed by western blotting.

### Statistical analysis

All experiments were performed with at least three independent biological replicates. Data were analyzed using GraphPad Prism 9. Comparisons among more than two groups were conducted using one-way analysis of variance (ANOVA) followed by Tukey’s post hoc test. Statistical significance was defined as p < 0.05.

## Acknowledgments

We thank Dr. Chih-Yen King for the yeast prion strain. This work was financially supported by the National Science and Technology Council (112-2311-B-002 −015 -MY3) to S.-C. Teng.

---

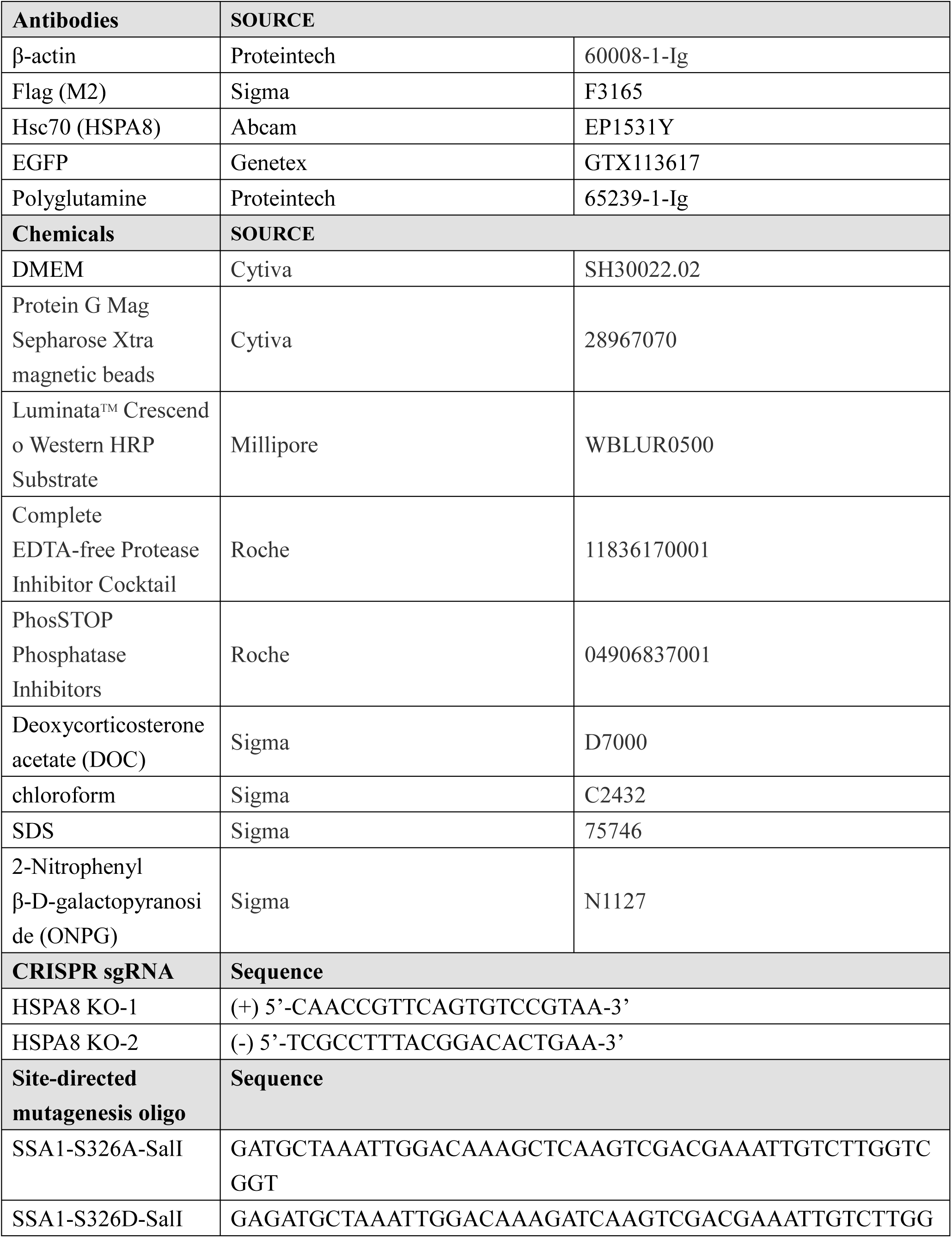
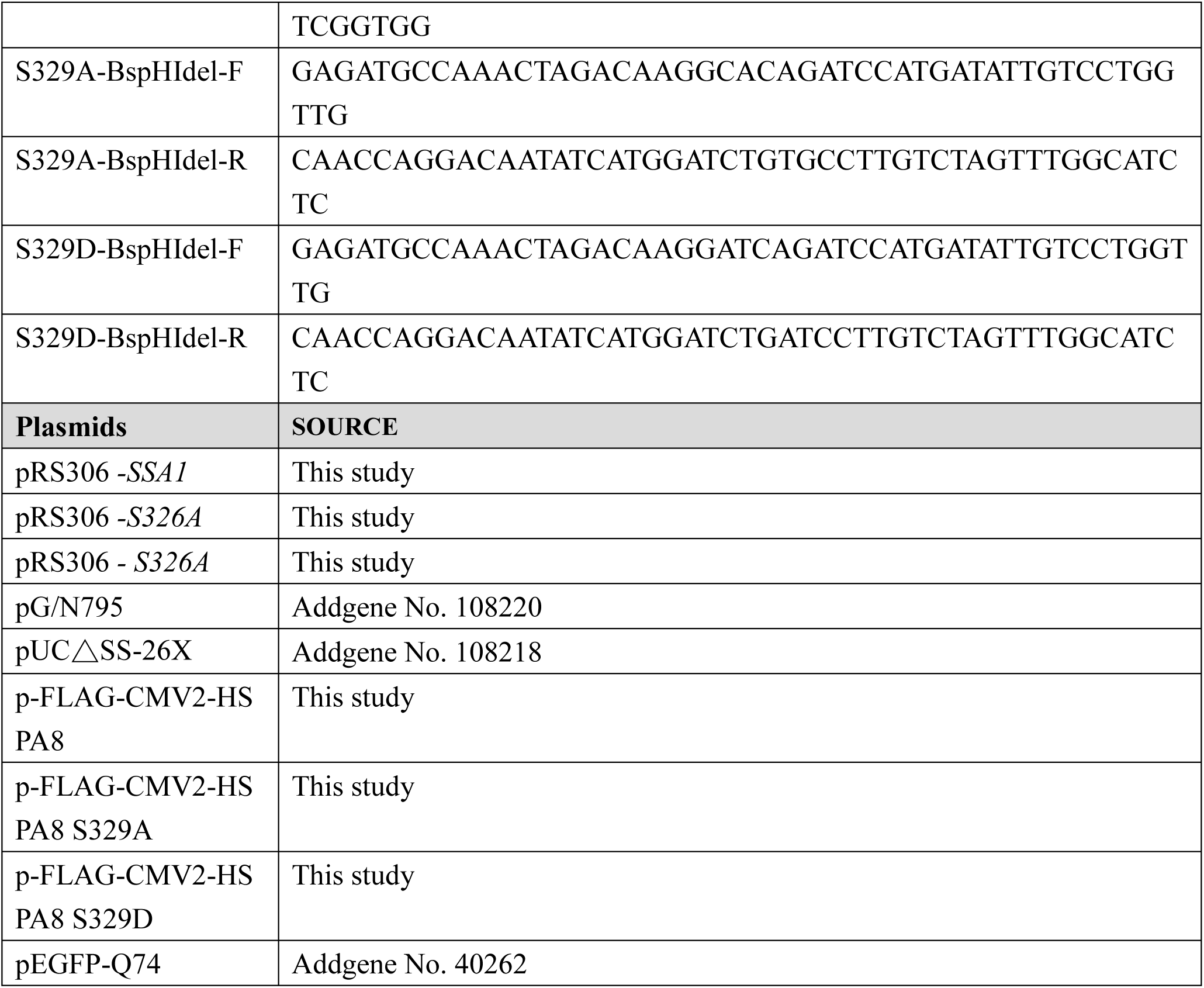
The Reagents and tools.

Yeast strains used in this study.

**Table. Yeast strains used in this study**
| Strain | Genotype |
| --- | --- |
| W303 | *MATa/α ura3-52/ura3-52 trp1 Δ2/ trp1 Δ2 leu2-3_112/ leu2-3_112 his3-11/ his3-11 ade2-1 can1-100/ ade2-1 can1-100* |
| CWY390 | W303 *ssa1::KanMX4* |
| CWY391 | W303 *ssa2:: HIS3MX6* |
| CWY392 | W303 *ssa1::KanMX4 ssa2:: HIS3MX6* |
| CWY429 | W303 *ssa1-S326A ssa2:: HIS3MX6* |
| CWY440 | W303 *ssa1-S326D ssa2:: HIS3MX6* |
| 5V-H19 | *SUQ5 ade2-1(UAA) can1-100 leu2-3,112 ura3-52* |
| CWY375 | 5V-H19 *ssa1::URA3* |
| CWY376 | 5V-H19 *ssa1-S326A* |
| CWY377 | 5V-H19 *ssa1-S326D* |

